# Korea4K: whole genome sequences of 4,157 Koreans with 107 phenotypes derived from extensive health check-ups

**DOI:** 10.1101/2022.12.25.521908

**Authors:** Sungwon Jeon, Hansol Choi, Yeonsu Jeon, Whan-Hyuk Choi, Hyunjoo Choi, Kyungwhan An, Hyojung Ryu, Jihun Bhak, Hyeonjae Lee, Yoonsung Kwon, Sukyeon Ha, Yeo Jin Kim, Asta Blazyte, Changjae Kim, Yeonkyung Kim, Younghui Kang, Yeong Ju Woo, Chanyoung Lee, Jeongwoo Seo, Dan Bolser, Orsolya Biro, Eun-Seok Shin, Byung Chul Kim, Seon-Young Kim, Ji-Hwan Park, Jongbum Jeon, Dooyoung Jung, Semin Lee, Jong Bhak

## Abstract

We present 4,157 whole-genome sequences (Korea4K) coupled with 107 health check-up parameters as the largest whole genomic resource of Koreans. Korea4K provides 45,537,252 variants and encompasses most of the common and rare variants in Koreans. We identified 1,356 new geno-phenotype associations which were not found by the previous Korea1K dataset. Phenomics analyses revealed 24 genetic correlations, 1,131 pleiotropic variants, and 127 causal relationships from Mendelian randomization. Moreover, the Korea4K imputation reference panel showed a superior imputation performance to Korea1K. Collectively, Korea4K provides the most extensive genomic and phenomic data resources for discovering clinically relevant novel genome-phenome associations in Koreans.

## Background

South Korea has perhaps one of the most extensive and convenient annual health check-up services. Every year, almost all Koreans aged over 40 receive a standardized health check-up, and the amount of individual clinical data is very extensive [1]. In 2020, we published 1,094 whole genomes with clinical information (Korea1K) by providing all the participants with an extensive and free standard health check-up showing the value of whole-genome data accompanied by clinical information mapping the genome diversity with practical applications [2]. Here, we present the second phase of the Korean Genome Project (KGP) with 4,157 sets of whole-genome data, Korea4K. It is accompanied by 107 types of clinical traits that have been donated by 2,685 healthy participants who acquired the health check-up reports from the hospitals of their choice in the past years. We manually annotated thousands of donated health reports that are matched with the whole-genome information. Therefore, apart from the increased number of samples, the main difference between Korea1K and Korea4K is that Korea4K’s clinical information is from very heterogeneous but fairly standard Korean health check-up centers, while Korea1K was from one very well-controlled university hospital health check-up center. This was also a testbed to assess how difficult it would be to merge data from the heterogeneous health check-up record system in a nation for a large-scale genome to phenome association analysis.

Previously, there were a few phenome-wide association studies (PheWASs) on Asian populations, but they were limited to chip or exome-based genotyping data. A Japanese PheWAS identified the genetic links among clinical traits, complex diseases, and cell-type specific patterns [3]. Another PheWAS using 10,000 Korean cohorts’ health check-up data from multiple lab sources showed network relationships between genes and phenotypes [4]. However, none of these studies covered the entirety of genomic variation, and they have limitations on genome-wide data analyses [5, 6].

A scientific contribution of this version of KGP is that we provide extensive genome-to-phenome association information with the largest genomic and clinical data from Korea to date to estimate how many samples and clinical parameters cover the whole genomic and common phenotypic diversity of Koreans. Korea4K contains 4,157 Korean genomes from East Asian ancestry, and 2,685 of them are accompanied by 107 types of clinical information such as height, waist circumference, weight, albumin/globulin ratio, basophil, direct bilirubin, low-density lipoprotein, high-density lipoprotein, mean corpuscular volume, and total cholesterol. The rest does not contain such kind of data because the biobank does not have phenotype information, or we were not able to collect it from the participants. Korea4K extends the efforts to completely map the totality of Korean genomic diversity, which can be a useful scope reference for disease risk prediction, diagnosis, and treatments in the future for personalized medicine.

As the second phase of the KGP, Korea4K not only extends the previously reported Korea1K [2] but also includes new multi-phenotypic association analyses, that is, analyses on markers that are associated with multiple phenotypes (pleiotropy), the genetic correlation between traits, and estimated causality relationship among traits through Mendelian randomization (MR) and 3D structure models for Korean specific missense variants. Combining these two omics data, we provide the community with the most extensive geno-phenotype association of healthy Korean participants. We have also applied the genomic variation data to the genotype imputation of low-frequency variants in the Korean population.

## Construction and content

### The largest Korean whole-genome variants data: Korea4K variome

A total of 64,301,272 single nucleotide variants (SNVs) and 8,776,608 Indels were called against the human genome reference (hg38) from the 4,157 Korean whole genomes, including 3,071 healthy controls (Table S1 and S2). It contains 3,063 newly added whole genomes sequenced by Illumina next-generation sequencing (NGS) platforms (HiSeq X10 and Novaseq 6000), in addition to the previous Korea1K dataset which was mostly generated by Illumina HiSeq X10. Using the variant data, we selected 3,617 samples with no kinship after initial sample filtering (see Methods). To exclude erroneous variants from sequencing batch effects from the heterogeneous Illumina NGS platforms and library preparation, we applied an allele balance bias measurement and finally acquired 12,713,580 erroneously called variant candidates (Fig. S1). After additional variant filtering (see Methods), we identified 45,537,252 variants including 42,124,137 SNVs, 36,029 double nucleotide variants (DNVs), 26,135 triple nucleotide variants (TNVs), 3,261,682 indels, and 89,269 other types of small variants from the 3,617 unrelated samples. We named this filtered Korean dataset the Korea4K variome (Fig. 1). A total of 23,689,147 variants were not present in the previous Korea1K variome. This unexpectedly large difference is likely derived from different batch effect filtering, and variant calling and filtering procedures, as well as new variants from the larger sample size. Consistent with the Korea1K study [2], most variants were located in intronic or intergenic regions and rarely in splicing sites or coding regions (Fig. S2), which is a sign of negative selection pressure in the population. Half of the total variants (21,941,879; 48.2%) were singleton or doubleton in the 3,617 unrelated samples, indicating that the Korean population’s genetic diversity is very low as the population diversity could be covered by fewer than 4,000 unrelated samples (Fig. 1a, Table S3). Almost all the common (allele frequency of > 0.01 and allele frequency of ≤ 0.05) and very common (allele frequency of > 0.05) variants were found to be already reported in the dbSNP database (99.70% and 99.97%, respectively), while more than half of the singleton and doubleton variants were newly discovered in this study (59.9% and 44.57%, respectively), indicating the new variant pool is well-exhausted in the Korean population by the 3,617 samples resulting in a large portion of individual specific novel variants in the Korean variome (Fig. 1a, Table S3). Only 3,092 and 3,569 unrelated individuals were needed to discover all the rare (allele frequency of > 0.001 and allele frequency of ≤ 0.01) and very rare (allele count of > 2 and allele frequency of ≤ 0.001) variants in the Korea4K variome, respectively (Fig. 1b) indicating that the Korea4K variome includes almost all the rare and very rare variants of Korean people of East Asian ancestry. It is notable that in our previous Korea1K data, the accumulated variant number curves did not reach a plateau [2]. Regarding common variants, only 481 and 161 unrelated individuals were necessary for common and very common variants, respectively, to cover the diversity which is close to the Korea1K statistics (440 and 132 samples). Essentially, the Korea4K variome statistics indicate the saturation of population diversity detection among Koreans. However, as expected, in the case of singleton and doubleton variants, the Korea4K variant discovery curve did not reach a plateau. This is due to each individual’s novel random variants and we will never reach a point of no novel variant discovery even with increased sample numbers.

**Fig. 1.**
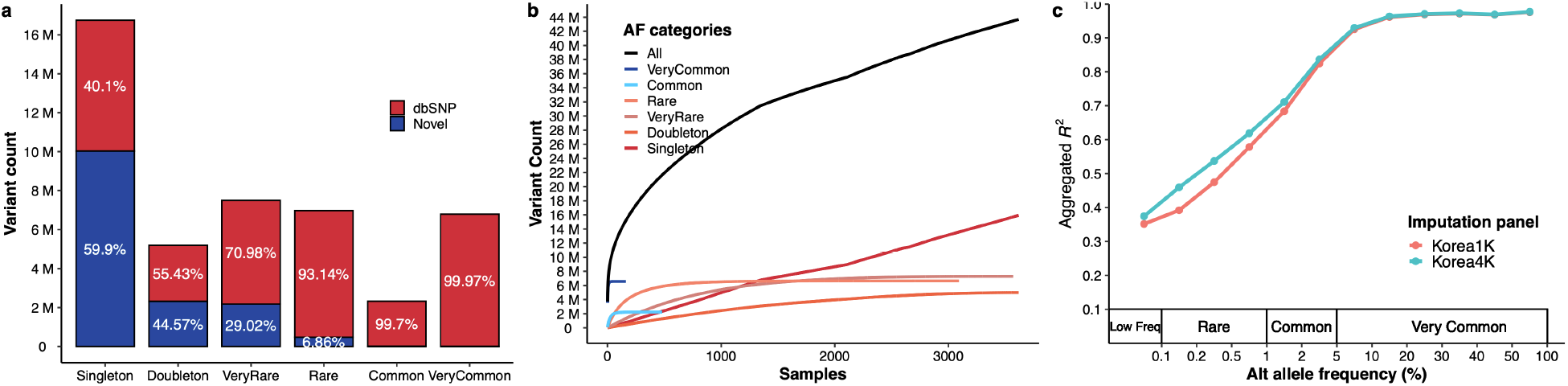
Korean variome profile and imputation evaluation using Korea4K. **(a)** The number of variants in the Korea4K variome is categorized by allele frequencies (AFs) among unrelated Korea4K genomes. Singleton, allele count = 1; doubleton, allele count = 2; very rare, allele count of > 2 and allele frequency of ≤ 0.001; rare, allele frequency of > 0.001 and allele frequency of ≤ 0.01; common, allele frequency of > 0.01 and allele frequency of ≤ 0.05; very common, allele frequency of > 0.05. **(b)** The number of discovered variants as a function of unrelated genomes. **(c)** Imputation performance evaluation using the Korea4K and Korea1K panels. The X-axis indicates alternative (Alt) allele frequency in the Korea4K variome. The Y-axis represents the aggregated *R*^2^ values of variants. We used variants that were overlapped by imputed results across two panels.

As a practical application, we constructed a Korea4K imputation reference panel from the 3,614 unrelated whole-genomes that showed a consistently better imputation performance than the Korea1K. The Korea4K panel was able to impute 198,805 more genotypes than the Korea1K panel (7,551,095 loci compared to 7,352,290) with the same dataset. Moreover, as expected, the Korea4K panel had better accuracy across all allele frequency categories than the Korea1K panel (Fig. 1c). The difference in aggregated *R*^2^ became larger for variants with allele frequency (AF) in Korea4K < 0.05 than for those in Korea1K, indicating higher accuracy in rare variants (Fig. 1c). In particular, the Korea4K imputation panel improved the imputation accuracy by 6% for the rare variants group compared to Korea1K on average.

As in Korea1K, the Korean population is genetically distinct from the Chinese and Japanese populations, confirmed by principal component analysis (PCA) with few outliers (Fig. 2a). We also found 62 missense variants out of 282,607 in Korea4K that had AFs significantly different from ten populations in the 1000 genome project (1KGP) from European Bioinformatics Institute (EBI), Cambridge, UK (Chi-squared test *P* < 5 × 10^-5^ against each of the ten populations, Extended data table S1). The genes containing such Korean-specific missense variants included *LILRB3, HLA-DRB5, IGLV5-48,* and *IGHV4-4* that are known to be associated with adaptive immunity, and *OR9G1* and *OR8U1* for olfactory receptors. Additionally, we found that twelve Korean-specific missense variants were in protein functional domains (Fig. 2d). Four of them were predicted to facilitate increased structural stability calculated in the protein 3D models built by AlphaFlod2 [7], while the other eight variants were predicted to cause decreased stability (Extended data table S2).

**Fig. 2.**
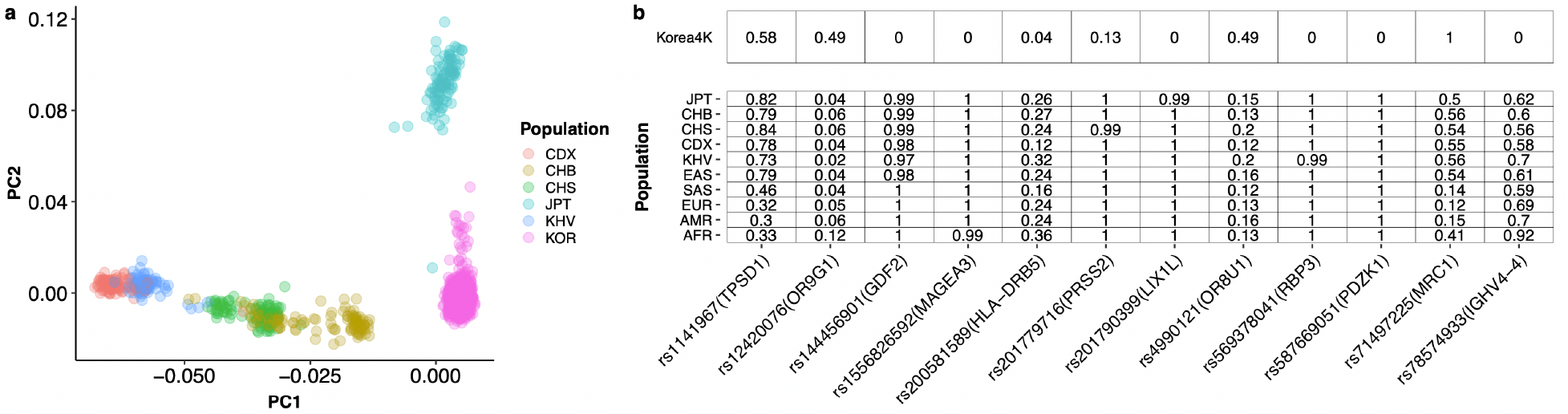
Comparison of Korea4K and 1KGP. **(a)** The results from principal component analysis of Korea4K and the 1KGP set of East Asian samples. **(b)** Allele frequency information of Korea4K and the populations in the 1KGP for the twelve Korean-specific missense variants located in protein functional domains. KOR: Korea4K; CDX: Dai Chinese; CHB: Han Chinese; CHS: Southern Han Chinese; PT: Japanese; KHV: Kinh Vietnamese; EAS: East Asians; SAS: South Asians; EUR: European; AMR: American; AFR: African.

### Whole-genome-wide association study (WGWAS)

Whole-genome-wide association studies (WGWASs) revealed that 2,324 variants from 157 unique loci had significant associations with 34 clinical traits from 37 WGWAS target traits *(P* < 5 x 10^8^; Fig. 3a-f, Extended data table S3). We used 90 clinical traits from the 107 phenotypes after filtering 27 traits with a high missing rate and biased distribution for WGWASs. Of the 90 traits, 54 were not confident in Quantile-Quantile plots and were excluded from further Mendelian randomization and pleiotropy analyses (see Methods). Among the 2,324 WGWAS significant variants, only 85 variants (31 loci) were reported in the GWAS catalog database [8]. The trait with the largest number of significantly associated loci was carbohydrate antigen 19-9 (CA19-9), a cancer antigen, with sixteen loci. Uric acid had the second highest number of significant loci with fourteen loci.

**Fig. 3.**
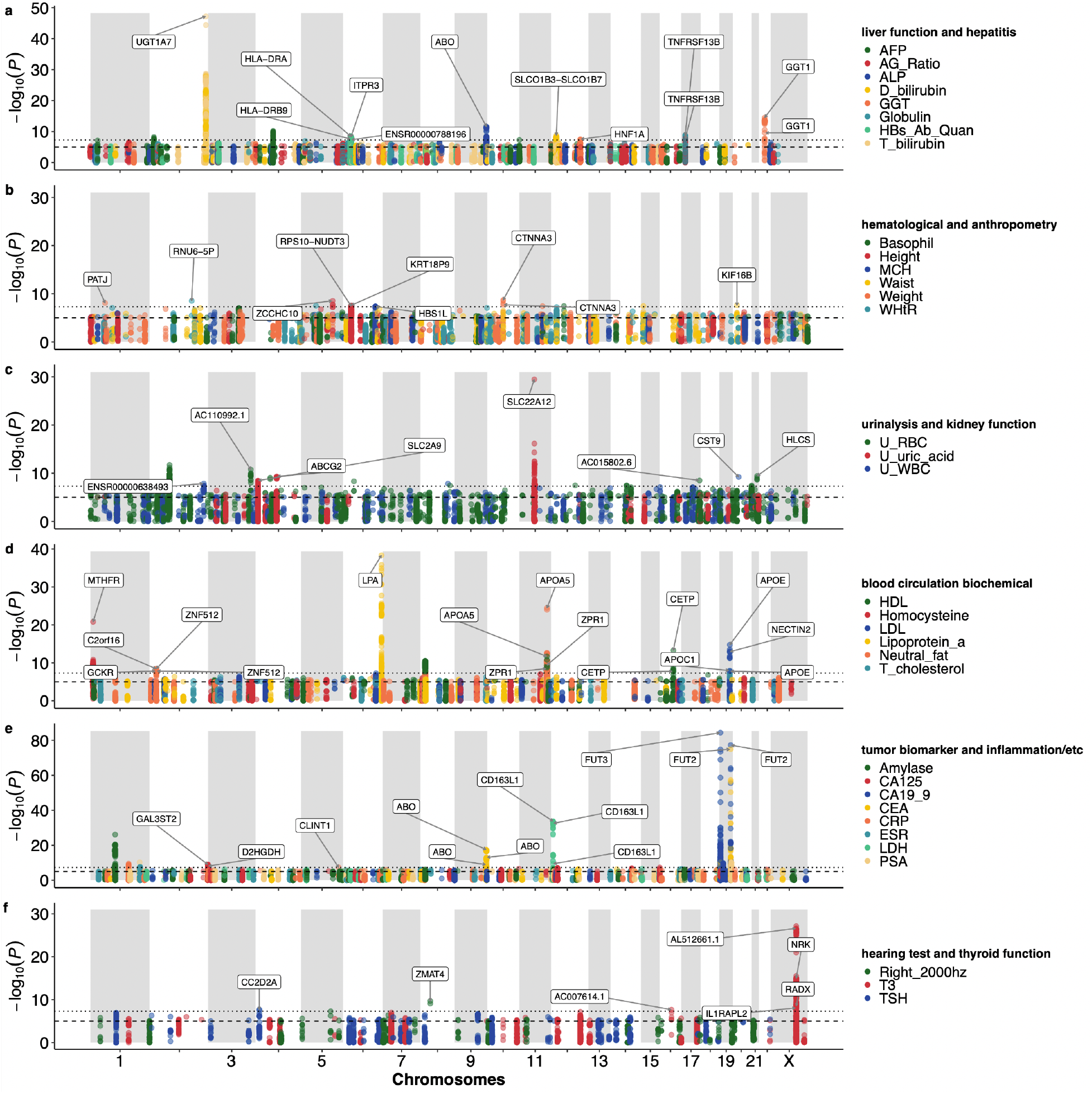
Whole-genome-wide association studies in Korea4K. **(a-f)** Whole-genome-wide association studies from 34 traits. Loci are presented only when index variants of the loci had significant *P*-value (*P* < 5 × 10^-8^) from the WGWAS. The dashed line indicates the suggestive threshold (*P* < 10^-5^). The dotted line indicates the significant threshold (*P* < 5 × 10^-8^).

Korea4K showed much stronger statistical power than the previous Korea1K study, identifying 1,356 new WGWAS variants (107 loci) from 28 common traits between Korea4K and Korea1K. Also, Korea4K had much lower (i.e., more significant) *P*-values than Korea1K for all the commonly found association variants between the two datasets (Fig. S3). Among the 107 loci containing the 1,356 new WGWAS variants, 798 Korea4K significant WGWAS variants from 73 loci had not been significant in Korea1K (Extended data table S3). Furthermore, twelve traits (albumin/globulin ratio, basophil, C-reactive protein, direct bilirubin, height, low-density lipoprotein, mean corpuscular volume, right hearing at 2000hz, thyroid stimulating hormone, total cholesterol, waist, weight) had 425 WGWAS variants that were significant uniquely in Korea4K, meaning no significant WGWAS variants from the twelve traits in Korea1K (Extended data table S3). For example, a missense variant, rs6431625 (*P* = 1.41 x 10^-23^), in *UGT1A3* was found to be associated with direct bilirubin in Korea4K. It was previously reported to be associated with circulating bilirubin levels [9]. Another Korea4K-specific missense variant is rs7412 (*P* = 2.86 x 10^-14^) in *APOE* which is associated with low-density lipoprotein (LDL) levels. Its association with cholesterol levels has been previously well-established [10]. Finding novel WGWAS variants in Korea4K was due to the increased sample size and subsequently increased variant number compared to Korea1K.

### Genetic correlation (GC) and phenotypic correlation (PC)

We found 27 traits with significant heritability among 89 quantitative traits (Fig. 4a; the lower boundary of genetic heritability > 0 with 95% confidence interval; Extended data table S4). A total of 24 pairs of traits showed a significant genetic correlation (FDR_GC_ < 0.05), measured as rG value, among 351 trait pairs between the 27 traits that showed significant heritability (Fig. 4, Extended data table S5). We found consistent results of Weight-Waist and body mass index (BMI)-Waist pairs, showing a significant genetic correlation in the UK Biobank data (http://www.nealelab.is/uk-biobank) with the same trend as our result (rG = 0.9, *P* = 10^-308^ in UK Biobank; rG = 0.9, *P* = 10^-308^ in UK Biobank, respectively). We identified 2,274 trait-trait relationships that had significant phenotypic correlation (FDR < 0.05, its 95% CI does not include 0) from trait-trait associations between 3,916 pairs of 89 quantitative traits (Fig. 4b, Extended data table S6). Most genetic and phenotypic correlations showed the same direction of correlation. The only two exceptions were waist/weight ratio (WWtR) – Urine white blood cell (U_WBC) and Waist-Creatine which showed opposite directions. This trend of Waist-Creatine has also been reported in a correlation database using UK-biobank data [11].

**Fig. 4.**
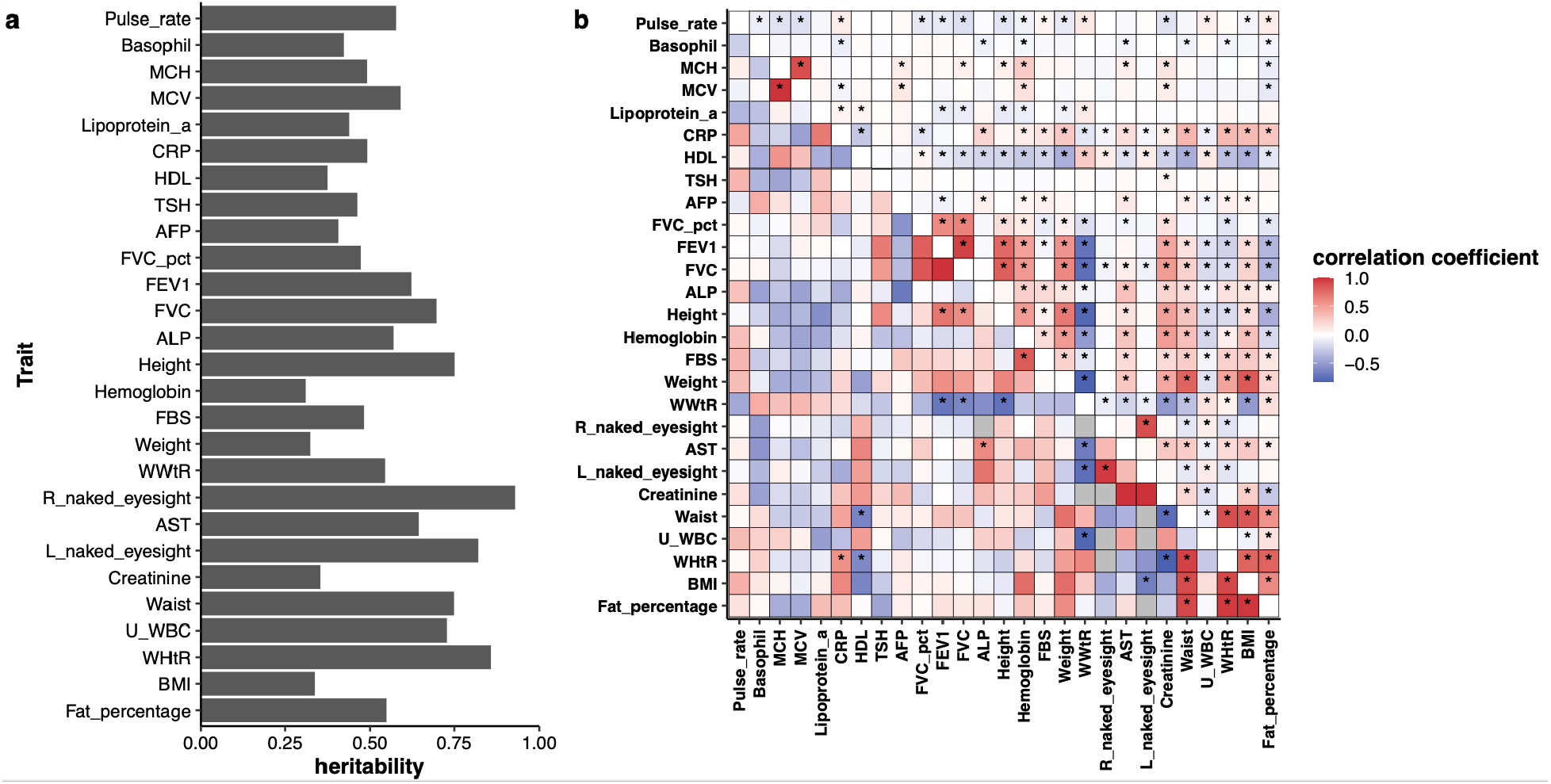
Genetic correlation and Phenotypic correlation in Korea4K. **(a)** Genetic heritability of 27 traits that showed at least a marginal significance. **(b)** Genetic correlation and phenotypic correlation between the 27 traits. The upper triangle indicates phenotypic correlation coefficient (Pearson’s) and lower triangle indicates genetic correlation coefficient (rG).

### Pleiotropy and Mendelian randomization (MR)

Out of the 37 WGWAS target traits, we detected 1,131 variants from 21 traits having suggestive associations (*P_GWAS_* < 10^-5^) with at least two traits, implying pleiotropic variants (Fig. 5, red edges; Extended data table S7). We devised the Variant-Sharing Index (VSI) to measure the degree of intersection between two phenotypes (Table 1). If the VSI equals zero, two traits share no suggestively associated variants, while 100 indicates the traits share all of them. The trait pairs with shared suggestive variants (SSVs) and the corresponding VSIs are listed in Table 1. Notably, we had only one variant, rs77913154 (chr5:18853857), that was shared among three traits: Globulin, AG_Ratio, and ESR (Extended data table S7). Interestingly, we found fifteen variants residing on *SOD2P1-AC095032.2-AC095032.1* locus forming pleiotropy between the serum amylase level and the level of CA125, a known ovarian cancer marker (VSI=2.3). Fourteen variants of the fifteen variants conform to the alteration of *AMY2B* level based on cis-eQTL results from GTEx Portal (ver.8), four of which were associated with expression in the pancreatic tissue. Already, there have been reports that patients with ovarian cancer manifest hyperamylasemia [12–14]. In terms of causality evaluation, a total of 127 trait pairs among 1,332 pairs of the 37 WGWAS traits were estimated to have significant causal relationships (FDR < 0.05, Fig. 5, Extended data table S8) from at least two of three different Mendelian randomization (MR) analysis methods (IVW: 166 pairs; MRPRESSO: 139; MR-Egger: 23). We found 59 unidirectional relationships and 68 bidirectional causal relationships (Extended data table S8).

**Fig. 5.**
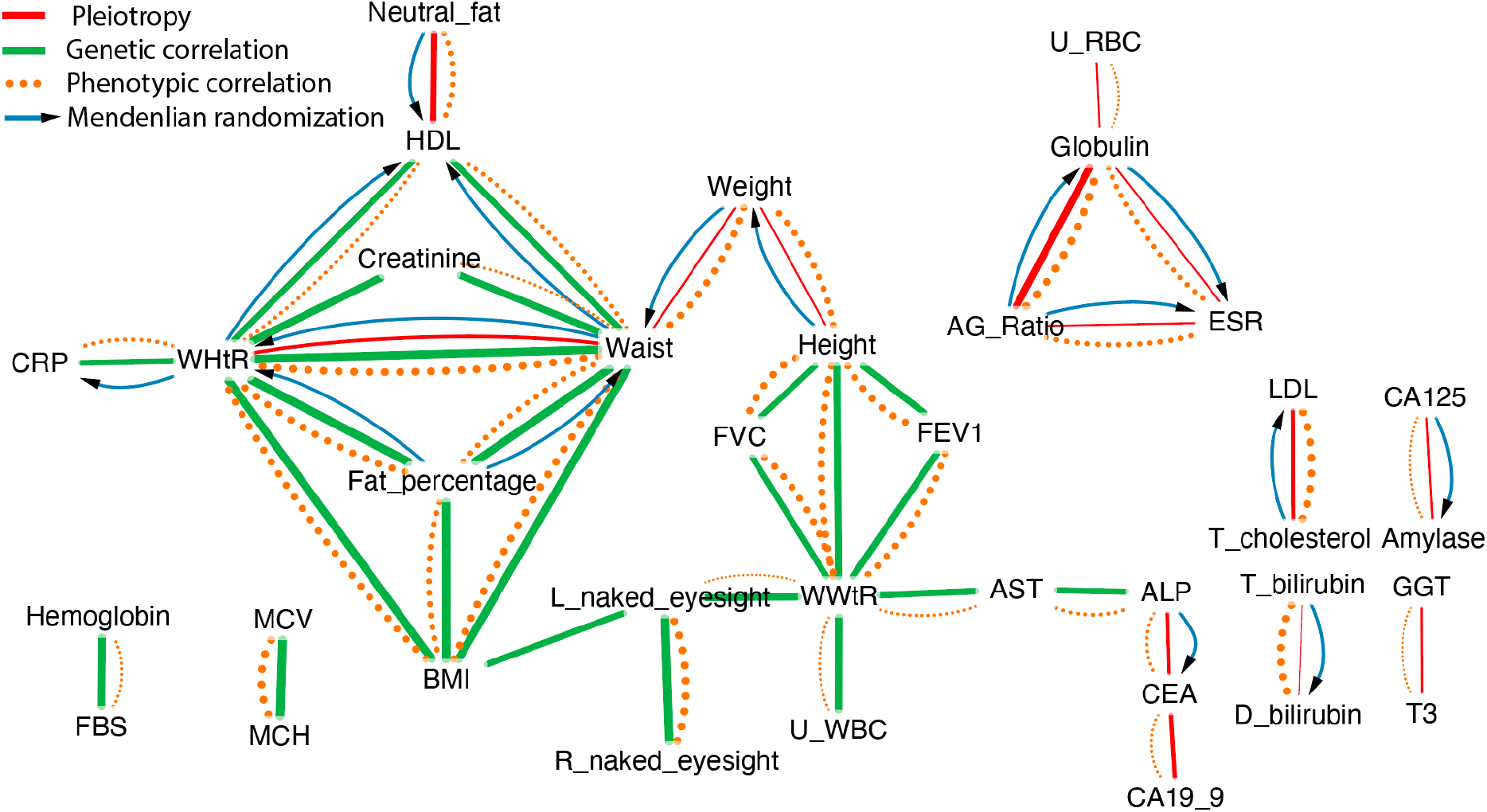
Graph visualization of genetic correlation, phenotypic correlation, pleiotropy, and Mendelian randomization. Green line indicates significant genetic correlation (GC), and the edge thickness indicates the absolute value of the correlation coefficient. Red line indicates trait pairs that have pleiotropic variants. Dotted orange lines indicate phenotypic correlation (PC), and the edge thickness indicates the absolute value of Pearson’s correlation coefficient. Blue arrow line indicates a causal relationship from Mendelian randomization (MR). MR and PC were visualized only when at least one of GC or Pleiotropy relationships was observed between the traits.

**Table 1.** Pleiotropic traits and Variant-Sharing Index (VSI)

| Trait1 | Trait2 | Suggestive variants in trait1 | Suggestive variants in trait2 | Shared variants | Total variants | VSI |
| --- | --- | --- | --- | --- | --- | --- |
| D bilirubin | T bilirubin | 638 | 632 | 569 | 701 | 81.2 |
| Globulin | AG_Ratio | 294 | 230 | 147 | 377 | 39 |
| HDL | Neutral fat | 348 | 398 | 191 | 555 | 34.4 |
| CEA | CA19_9 | 221 | 264 | 74 | 411 | 18 |
| T cholesterol | LDL | 74 | 238 | 38 | 274 | 13.9 |
| WHtR | Waist | 177 | 100 | 31 | 246 | 12.6 |
| ALP | CEA | 153 | 221 | 35 | 339 | 10.3 |
| T3 | GGT | 542 | 125 | 23 | 644 | 3.6 |
| CA125 | Amylase | 202 | 466 | 15 | 653 | 2.3 |
| Weight | Waist | 123 | 100 | 5 | 218 | 2.3 |
| Height | Weight | 173 | 123 | 2 | 294 | 0.7 |
| ESR | AG_Ratio | 163 | 230 | 1 | 392 | 0.3 |
| Globulin | ESR | 294 | 163 | 1 | 456 | 0.2 |
| U_RBC | Globulin | 627 | 294 | 1 | 920 | 0.1 |

### Summary results of the four phenomics analyses

We summarized the four phenomics analyses (Genetic correlation, Phenotypic correlation, Mendelian randomization, and pleiotropy) by visualizing them in network plots (Fig. 5). In general, the discovered trait-trait pairs of genetic correlation, Mendelian randomization, and pleiotropy analysis results were not often overlapping. Nevertheless, the network visualization suggests distinguishable association patterns of the two secondary body measures, WHtR and WWtR with other phenotypes. WHtR had associations with the C-reactive protein (CRP), creatine, and HDL. On the other hand, WWtR was associated with aspartate aminotransferase (AST), forced expiratory volume (FEV1), forced vital capacity (FVC), and urine white blood cell (U_WBC). Despite their similarity, they may reflect different biological mechanisms.

Genetic correlation and pleiotropy are found exclusive of each other, despite both measures having shared genetic components of two different traits. GC is mainly observed from body measures such as waist, weight, height, and Left-naked eyesight. Pleiotropy was more on the relationship between metabolites in blood such as LDL, bilirubin, or CEA. The only overlap is WHtR-Waist, where one is derived from the other. MR analysis suggests a causal relationship between phenotypic correlations. For example, the Fat-percentage influences Waist and WHtR, which is followed by the influence on HDL and CRP. The result is concordant with previous reports that body fat percentage and CRP are correlated [15, 16]. ALP and CEA showed potential causality, as well as the shared variants between them (pleiotropy near *ABO* gene). Many previous studies reported them together as targets for diagnosing cancer and monitoring metastasis [17–19]. Nevertheless, their molecular-level relationship has not been suggested. Our phenomics results also suggested that Waist/Height ratio is a linked trait in the association of paired traits, and the association is probably derived from indirect causation.

### Sample collection and whole-genome sequencing

We collected 2,848 blood samples or already processed DNA samples from Korean individuals. A total of 1,094 whole-genome sequencing (WGS) datasets originating from our previous study (Korea1K) and 215 WGS data from publicly available Clinical & Omics Data Archive (CODA) were added to the aforementioned dataset [2]. The genomic DNA was extracted using the DNeasy Blood & Tissue kit (Qiagen) from whole blood samples. We constructed the whole-genome sequencing library from the DNA by using the TruSeq Nano DNA Sample Prep kit (Illumina) kit. Whole-genome sequences of the 2,848 samples were generated by the Illumina Nova-seq 6000 platform. Sample collection and sequencing were approved by the Institutional Review Board (IRB) of the Ulsan National Institute of Science and Technology (UNISTIRB-15-19-A and UNISTIRB-16-13-C).

### Joint genotype calling

Adapter contamination was trimmed using Cutadapt (ver. 1.9.1) [20] with a forward adapter (‘GATCGGAAGAGCACACGTCTGAACTCCAGTCAC’) and reverse adapter (‘GATCGGAAGAGCGTCGTGTAGGGAAAGAGTGT’) and with a minimum read length of 50 bp after trimming. We mapped the whole-genome sequencing reads from 4,157 samples to the human reference genome (hg38) using BWA-mem (ver. 0.7.17) with the ‘-M’ option and alt-aware mode [21]. The mapped reads were sorted by genomic coordination using Picard (ver. 2.20.3). We marked the PCR-duplicates and recalibrated the base quality of the mapped reads using the MarkDuplicates and BaseRecalibrator module in Picard (ver. 2.20.3), respectively. A total of 3,156 samples had a mapping depth of ≥ 20 × (Fig. S20). Individual genotypes were called in GVCF format by HaplotypeCaller in GATK (ver. 4.1.3) with ‘--genotyping-mode DISCOVERY -standcall-conf 30 -ERC GVCF’ options [22]. We merged the individual genotypes to a single GVCF for each chromosome using CombineGVCFs in GATK (ver. 4.1.3) [22]. We jointly genotyped the merged GVCF with the genotypeGVCF module in GATK (ver.4.1.3) [22]. Variant quality of the joint genotypes was recalibrated using the VQSR module in GATK (ver. 4.1.3) [22].

### Sample and variant filtering

After joint genotyping, we filtered out a total of 540 participants on the criteria that are listed below using SelectVariants in GATK (ver. 4.1.3) with ‘--remove-unused-alternates’ option to remove unused variants [22].

1. showing high genotype missing rate (>10%): nine samples
2. having too high or low heterozygous variants ratio compared to homozygous variants per sample (3 s.d.): four samples
3. having relatedness to other samples: 428 samples
4. having non-Korean genetic background from PCA analysis with 1KGP set: seven samples
5. reported to have a rare disease: 40 samples
6. 52 samples who became not applicable for this study

Finally, the Korea4K variome data included 3,617 participants’ genomes. To detect variants which were probably called because of a sequencing batch effect, we measured average allele balance of the alleles. Then, we excluded 12,713,580 variants that had average allele balance of the loci out of the range of ± 1 × standard deviation (SD) from a genome-wide average of allele balance to remove the sequencing batch effect (Fig. S1). We also excluded the variants which had a genotyping rate of < 0.9 for downstream variant analysis. The variants in the final variome set were annotated using Variant Effect Predictor (VEP) with Ensemble database (ver. 101) [23].

### Principal Component Analysis (PCA) with the EBI’s 1KGP genome data

The interpopulation genomic structure was evaluated by projecting the first two PCs determined via PCA of SNVs from both Korea4K and East Asian populations from 1KGP. We merged variants from the Korea4K and 1KGP sets and then filtered out variants with the following criteria: (i) biallelic SNVs with a MAF < 1%; (ii) biallelic SNVs with an HWE *P* < 10^-6^; (iii) biallelic SNVs with a missing genotype rate of > 0.01. Extracted variants were LD pruned using “ --indep 200 4 0.1” option in PLINK (ver. 1.90b3n) [24], yielding 330,350 sites. PCA was carried out using PLINK (ver. 1.90b3n) [24].

### Korean-specific missense variants

We collected allele frequency data from ten populations (African (AFR), American (AMR), European (EUR), South Asian (SAS), East Asian (EAS), Japanese in Tokyo (JPT), Kinh Vietnamese (KHV), Han Chinese in Beijing (CHB), Han Chinese Southern (CHS), and Chinese Dai in Xishuangbanna (CDX)) from EBI’s 1KGP database [25]. For each Korea4K variant, we compared its allele frequency to the allele frequency of all of the ten populations using the Chi-squared test. We selected variants that were specific to the Korean when the *P*-value of the Chi-squared test to the ten populations was less than 5 × 10^-5^.

### Protein structure modeling and thermodynamic stability measurement

We constructed the mutant-type (MT) protein sequences of the Korean-specific missense variants by substituting the reference protein sequences found in the Ensembl database (ver. 101) [26]. We modeled the structures of the wild-type (WT) and mutant-type protein models using AlphaFold2 (ver. 2.0) with the ‘--max_template_data 2022-05-09 --db_preset reduced_dbs’ option with default databases downloaded by AlphaFold2 [7]. We used the InterPro database [27] to determine whether a missense variant was located in the domain region within the protein sequence. We extracted the domain region from the WT and MT protein 3D models and excluded domains that had less than 50 amino acids. Afterwards, we calculated ΔG_WT_ and ΔG_MT_ using the ‘Stability’ command of foldX [28] to measure the protein thermodynamic stability. Finally, we measured the change in protein thermodynamic stability between the two models by calculating the difference between the WT and MT domain models (ΔΔG = ΔG_MT_ – ΔG_WT_).

### Imputation

We constructed an imputation reference panel of Korea4K and Korea1K sets which includes 3,614, and 873 Korean individuals, respectively. A total of 26,210,741 and 15,649,303 autosomal biallelic variants with a missing genotype call rate of < 0.1 and minor allele count > 1 (not a singleton) were extracted for the Korea4K and Korea1K panel, respectively. The extracted variomes were phased into haplotype using SHAPEIT2 (ver. v2.r904) [29]. We used the same test dataset as in the previous study [2]. The phased test data was imputed using the imputation reference panel by Minimac3 (ver. 2.0.1) [30]. We estimated imputation accuracies using squared Pearson’s correlation coefficients (*R*^2^) between the true genotypes and imputed genotype dosages.

### Clinical information

We collected or calculated 107 clinical parameters (93 quantitative and 14 qualitative traits; Extended data table S9) along with genome data from 2,685 samples among the Korea4K samples. A total of 3,383 clinical datasets (including multiple time points per sample) from regular health checkups carried out by various hospitals and clinics throughout Korea were collected from 2,685 participants between 2016 and 2019. When a single participant had multiple clinical datasets, the most recent one was chosen for the following analysis. Four quantitative clinical traits and 12 qualitative traits were excluded from the further analysis, since the traits were missing from more than 90% of participants due to health check-up reports heterogeneity, or the traits that were qualitative and biased to one category (more than 1:4). Standard Weight was also removed from the analysis, because the trait was not an inherently correct representation of the sample’s clinical data but rather a recommended value. Three traits (Hepatitis B virus antibody, antigen, and hepatitis C antibody) contained both quantitative and qualitative values. Therefore, both of the values were utilized for analysis, i.e, Hbs_Ab_Quan and Hbs_Ab_Binary. Phenotypic correlations were calculated by Pearson’s method.

### Whole genome-wide association study (WGWAS)

SNVs and indels with a MAF <1%, HWE *P* < 10 ^−6^, and a missing genotype rate of > 0.01 were excluded from the analysis using PLINK (ver. 1.90b3n) [24]. A total of 90 WGWAS (88 quantitative and 2 qualitative traits) were performed with a total of 3,617 individuals and 7,782,381 variants. Each WGWAS had a different number of individuals that included those who had the target clinical traits. The WGWAS was performed using linear and logistic regression under an additive genetic model with PLINK (ver. 2.00 alpha) [31] for quantitative and qualitative traits, respectively. Sex, age, age^2^ (age squared), body mass index (BMI), and the top ten principal components of SNV genotypes were included in the model as covariates. BMI was excluded from covariates in the WGWAS for BMI itself and degree of obesity. We rejected 53 traits from further analysis based on QQ-plot analysis (Fig. S4-19). We used 5 × 10^-8^ for a whole-genome-wide significance threshold. The 7,782,381 variants were clumped into 466,938 loci based on linkage disequilibrium (LD) information using PLINK (ver. 1.90b3n) with ‘--clump-p1 1, --clump-p2 1, --clump-r2 0.1, --clump-kb 250, and --clump-index-first’ options [24].

### Measuring heritability and genetic correlation

We calculated genetic relatedness among individuals from SNPs by genetic relationship matrix (GRM) in genome-wide complex trait analysis (GCTA) (ver. 1.93.2) with ‘ --autosome --maf 0.01 --make-grm’ options [32]. We estimated the genetic heritability of 87 quantitative traits using GCTA (ver. 1.93.2) with ‘--reml --grm’ options [32]. We estimated the genetic correlations (GC) using the bivariate genome-based restricted maximum likelihood (GREML) algorithm [33] in the GCTA (ver. 1.93.2) with ‘--reml-bivar --grm --reml-bivar-lrt-rg’ options [32]. Two of the 253 trait pairs were excluded since the log-likelihood did not converge.

### Calculation of Variant Sharing Index (VSI)

The variant sharing index (VSI) is a Jaccard score to measure how many pleiotropic components exist out of all significant variants from *i*-th and *j*-th traits, which is defined as

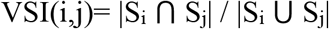

where S_i_ and S_j_ denote sets of significant variants for the *i*-th and *j*-th traits, respectively. The VSI increases as two traits have more pleiotropic variants among their significant variants.

### Pleiotropic variants with tissue-specific expression regulatory function

We annotated the gene symbol of the pleiotropic variant by using Ensemble database (ver. 101) [26]. In case of intergenic variants, we annotated the genes which were located the nearest in both directions of the variant. The single tissue eQTL data (ver. 8) from the GTEx portal were used to investigate the eQTL of pleiotropic variants in Korea4K.

### Investigation of potential causal relationships between traits based on Mendelian randomization (MR)

We used the Mendelian randomization method to investigate potential causal relationships among 1,332 combinations of an exposure trait and an outcome trait among 37 clinical traits. MR is computed from the linear regression analysis between the effects of SNPs on an exposure trait and their effects on an outcome trait. We chose the SNPs with suggestive WGWAS results (*P*-value < 10^-5^) with exposure traits as the instrument variables. In case multiple SNPs existed in the LD block, the one with the smallest *P*-value was chosen. We rejected 40 SNPs, which were detected as outliers of linear regression from MR-PRESSO software (1.0) [34] with ‘NbDistribution=10000 and SignifThreshold=0.05’ options, from further analysis. MR coefficients were computed using the chosen SNPs by three different methods: the Inverse-variance weighted (IVW) and MR-Egger method of TwoSampleMR package (v.0.5.6) [35] and MR-PRESSO software (1.0) [34]. Finally, we selected 36 significant causal relationships that overlapped at least two of three methods (IVW, MR_Egger, and MRPRESSO). All analyses were performed with default options.

### Utility and discussion

Batch effect exacerbated by sequencing platform and library preparation bias is a critical problem in very large population genome association studies, especially with clinical data from heterogeneous health check-up centers. In the future, more and more diverse whole-genome data with extensive clinical data will be publicly available, and it is inevitable that they will be merged for more precise whole genome-to-phenome association research. Korea4K is not an exception in that regard, and in one homogeneous population WGWAS, it was necessary to consider and factor in a great deal of sequencing and clinical data batch effects and errors. We attempted to minimize the errors by using allele balance with optimal filtering criteria and time-consuming manual checks on health reports that were donated by the participants. The largest challenge of Korea4K project was cleaning up heterogeneous clinical data from different health check-up centers. Another major issue was that the health check-up data heterogeneity caused reduced numbers of participants’ common traits with which to compare. Some of the health data were from past years’ health checkups from heterogeneous hospitals throughout Korea. This heterogeneity in location and time was not an intentional experimental design but was in order to reduce the cost of performing expensive one-center health check-ups for the Korea4K participants. Therefore, WGWAS along with standardized and unified national and public health check-up data will greatly benefit future wholegenome-wide association studies.

Although 4,157 seems like a large number, we found the sample size in this study was still not large enough to detect weak association signals. The Korea4K variome with matched phenotype information has allowed us to estimate genomic correlation across various phenotypes using GREML [32]. GREML has been reported to have higher accuracy compared to methods, such as linkage disequilibrium score regression (LDSC), using only summary statistics from GWAS [36]. For example, the minimum heritability score was 0.34 (Degree of obesity) among the traits detected as statistically significant. The statistical power of our maximum 2,685 subjects and FDR < 0.05 is estimated to be 0.72 for detecting traits with heritability of 0.3 or higher (Calculated from GCTA-GREML Power Calculator) [37]. This will increase to 0.97 with 4,000 subjects.

WGWAS, whole-genome-wide association, not chip-based GWAS, performs better in geno-phenotype association studies, and we suggest WGS for future studies for its genetic data completeness. Our pleiotropy analysis based on the WGWAS made it possible to reveal the portions of genetic association across multiple traits. For example, we could identify the variants in the well-known pleiotropic relationships such as ALP-CEA by *ABO* locus (35 variants), Neutral_Fat-HDL by *LPL* locus (181 variants) and Total cholesterol-LDL by *TOMM40,* and *APOE* locus (4 and 2 variants, respectively). These loci and their corresponding trait pairs were previously reported from chip-based GWAS summary results [38, 39]. However, we found more pleiotropic variants thanks to whole-genome-wide, unbiased coverage of WGS. Notably, the four methods that we adopted produced discrete trait-trait relationships, which means that multiple phenomics methods should be applied to investigate specific relationships and mechanisms among clinical traits or diseases. In other words, phenomics analyses were limited and not powerful enough to discover novel and indirect associations with current datasets.

One of the purposes of Korea4K was to build a reference dataset to discover unknown whole-genome to phenome associations that can be detected from samples of healthy people. This, however, is contradictory and it limited us in discovering clear pathogenic associations because most of the participants examined in WGWAS were healthy without any severe aberrant phenotypes or diseases that could bring us clues for interesting omics analyses.

There are three important limitations of our study. The first is we failed to acquire long DNA sequencing reads from the healthy participants for building a structural variation reference set for the Korean population. The second is the lack of epigenomic data from the 4,157 samples. This was mostly due to high costs for generation and computing long-read based assemblies and sequencing methylated DNA sites. The third one, which is perhaps the most relevant for the purpose of performing association studies for healthcare is that we failed to acquire more rare and severe disease data from patients, accompanied by precise clinical and multiomics data. We have excluded a small number of rare disease cases, as those required a large amount of sequencing data from genome, transcriptome, and methylome to perform precise functional analyses. Large-scale pathological whole-genome-wide omics data will become a powerful set for genome-phenome level association studies to detect causal markers for the prediction and diagnosis of health conditions in future studies.

## Conclusions

In conclusion, we reported the Korea4K dataset, the second phase of the KGP, which includes a large-scale Korean variome database as well as information on clinical traits. We believe Korea4K is a valuable genome-phenome resource for discovering clinically relevant novel associations and can be contributed to various fields of genomics research as a Korean reference panel.

## Abbreviations

KGP: Korean Genome Project
SNV: single nucleotide variant
NGS: Next-generation sequencing
DNV: double nucleotide variant
TNV: triple nucleotide variant
AF: allele frequency
AC: allele count
PCA: principal component analysis
1KGP: the 1000 genome project
WGWAS: Whole-genome-wide association study
GC: genetic correlation
PC: phenotypic correlation
MR: Mendelian randomization

## Declarations

### Ethics approval and consent to participate

Sample collection and sequencing were approved by the Institutional Review Board (IRB) of the Ulsan National Institute of Science and Technology (UNISTIRB-15-19-A and UNISTIRB-16-13-C). The biospecimens for this study were provided by Ulsan Medical Center and the Biobanks of Gyeongsang National University Hospital, Chungbuk National University Hospital (18-27, 20-04), and Kyung Hee University Hospital (2018-4, 2019-4, 2019-6), the members of the National Biobank of Korea, which is supported by the Ministry of Health, Welfare and Family Affairs. 215 whole-genome-seq data used in this study were provided by the Clinical & Omics Data Archive (CODA; http://coda.nih.go.kr), CODA accession number S000680. Written informed consent was acquired from all other subjects.

### Availability of data and materials

Allele frequency information of variants is publicly available under http://koreangenome.org. Raw sequencing data, individual genotype information, and clinical trait data will be as easily and freely available as possible upon request and after approval from the Korean Genomics Center’s review board in UNIST. Information about the Korean Genome Project and other data sharing can be found at http://koreangenome.org.

### Author contributions

S.J., Hansol C., Y. J., W.C. and Jong B. wrote the manuscript. S.J., Hansol C., Y. J., Hyunjoo C., K.A., H. R., Jihun. B., H. L., Yoonsung. K., S. H., C.L., and J. S. conducted the data analysis. C. K., Yeonkyung K., Younghui K., and Y. J. W. performed wet-lab experiments. S. J., Yeo Jin K., B. C. K., S.L., and Jong B. designed the study. S.J., Hansol C., Y.J. W.C., A.B., D. B., O. B., E. S., S. K., J. P., J. J., D. J., S. L., and Jong B. revised the manuscript. S. L. and Jong B. jointly supervised the study.

## Acknowledgments

We appreciate all participants and Ulsan citizens who supported this project. We also thank Ju Yeon Park, Sangryoul Han, Jungae Shim, Nayoung Kim, Seung Gu Park, Byoung-Chul Kim, Jungeun Kim, Neung-hwa Park, Suan Cho, and Yeshin Park for supporting this project. This work was supported by Biodatafarm computing infrastructure funded by the Ulsan metropolitan city government. We thank the Korea Institute of Science and Technology Information (KISTI) for providing us with the Korea Research Environment Open NETwork (KREONET). We thank our collaborators in NCSRD, KRISS, and C.-G. Kim. The Genotype-Tissue Expression (GTEx) Project was supported by the Common Fund of the Office of the Director of the National Institutes of Health, and by NCI, NHGRI, NHLBI, NIDA, NIMH, and NINDS. The data used for the analyses described in this manuscript were obtained from the GTEx Portal on 11/25/2021. We thank Jaesu Bhak and Maryana Bhak for editing the manuscript.

## Supplementary information

**Additional file 1: Supplementary materials; Fig. S1** Variants batch effect of DNA sequences. **Fig. S2** Variants distribution based on variant location and allele frequency category in Korea4K. **Fig. S3** Power comparison of whole-genome-wide association study between Korea4K and Korea1K. **Fig. S4** QQplots for the whole-genome-wide association tests of the traits on the anthropometry category. **Fig. S5** QQplots for the whole-genome-wide association tests of the traits on blood circulation biochemical category. **Fig. S6** QQplots for the whole-genome-wide association tests of the traits on blood circulation physics category. **Fig. S7** QQplots for the wholegenome-wide association tests of the traits on diabetes category. **Fig. S8** QQplots for the wholegenome-wide association tests of the traits on electrolyte category. **Fig. S9** QQplots for the whole-genome-wide association tests of the traits on hearing test category. **Fig. S10** QQplots for the whole-genome-wide association tests of the traits on hematological category. **Fig. S11** QQplots for the whole-genome-wide association tests of the traits on hepatitis category. **Fig. S12** QQplots for the whole-genome-wide association tests of the traits on inflammation and etc category. **Fig. S13** QQplots for the whole-genome-wide association tests of the traits on kidney function category. **Fig. S14** QQplots for the whole-genome-wide association tests of the traits on liver function category. **Fig. S15** QQplots for the whole-genome-wide association tests of the traits on pulmonary function category. **Fig. S16** QQplots for the whole-genome-wide association tests of the traits on thyroid function category. **Fig. S17** QQplots for the whole-genome-wide association tests of the traits on tumor biomarker category. **Fig. S18** QQplots for the whole-genome-wide association tests of the traits on urinalysis category. **Fig. S19** QQplots for the whole-genome-wide association tests of the traits on vision category. **Fig. S20** Mapping depth distribution of Korea4K genomes. **Table S1** Sample count in Korea4K. **Table S2** Variant count in Korea4K before sample and variant filtering. **Table S3** Variant count based on variant categories and reported to dbSNP.

**Additional file 2: Extended data tables. Extended data table S1** Allele frequency information of populations for 62 Korean-specific missense variants. **Extended data table S2** Prediction of changes in protein thermodynamic stability according to missense variant. **Extended data table S3** List of the GWAS variants which have association significance *P*< 5E-8. **Extended data table S4** Genetic heritability measurement. **Extended data table S5** Genetic correlation measurement. **Extended data table S6** Phenotypic correlation estimation. **Extended data table S7** Pleiotropic variants. **Extended data table S8** Mendelian randomization results. **Extended data table S9** Statistics of clinical information.

